# An efficient protocol to derive primordial germ cell like cells *in vitro* from murine embryonic stem cells

**DOI:** 10.1101/2025.05.12.653613

**Authors:** Umesh Kumar, Nithyapriya Kumar, P Chandra Sekhar, Kumarasamy Thangaraj

## Abstract

Primordial germ cells (PGCs) are the precursors of germ cells, specified from a subset of epiblast stem cells at the post-implantation stage of mammalian embryonic development. Over the past decade, primordial germ cell-like cells (PGCLCs) have been successfully derived *in vitro* from mouse and human embryonic stem cells (ESCs). These PGCLCs can further differentiate into mature sperm upon transplantation into neonatal testes. Such advancements offer promising strategies to address male infertility, which affects approximately 15% of couples worldwide. The discovery of induced pluripotent stem cells (iPSCs), which possess similar potential to ESCs, further raises hope for patient-specific therapeutic interventions. However, a major challenge remains: the low efficiency of *in vitro* PGCLC differentiation. To overcome this, we developed an optimized PGCLC differentiation strategy using a Dppa3-mCherry knock-in reporter mouse ESC line, TDM11. By testing various culture conditions, we identified ascorbic acid as a potent inducer of PGCLC differentiation from ESCs, providing a potential avenue for enhancing *in vitro* germ cell derivation.

## Introduction

Germ cells are the precursors of gametes, responsible for genome recombination and genetic transmission to the next generation. In mammals, germ cell development begins with unipotent primordial germ cells (PGCs), which undergo sequential differentiation to form sperm in males and ova in females. Fertilization of these gametes results in a totipotent zygote, whose totipotency gradually diminishes during the transition from blastomere to blastocyst. In mice and humans (and presumably all mammals), germ cell lineage specification occurs in the early embryo through epigenesis, where a subset of epiblast cells is induced by paracrine signaling from surrounding tissues (de Sousa Lopes et al., 2007; Kurimoto et al., 2008). While major molecular events of germ cell development are well characterized, the regulatory mechanisms governing this process remain poorly understood. Infertility affects approximately one in seven couples globally, with idiopathic molecular causes accounting for about 60% of cases (Sharlip et al., 2002). This underscores the importance of identifying molecular regulators that control key events in germ cell development.

A major challenge in studying human germ cell development is the limited availability of early embryonic tissue. However, recent advancements have enabled the *in vitro* derivation of PGCs and mature germ cells (sperm and oocytes) from embryonic stem cells (ESCs) (Hayashi et al., 2011; Hikabe et al., 2016; Yamashiro et al., 2020). These breakthroughs provide a platform for investigating the molecular mechanisms of germ cell development *in vitro* (Irie et al., 2015; Smela et al., 2019; Sugawa et al., 2015) and offer potential applications in regenerative medicine to restore fertility (Sosa et al., 2018). Despite these advancements, *in vitro* germ cell differentiation remains challenging due to its complexity, inefficiency, and the heterogeneity of developmental stages. These limitations hinder both the functional characterization of novel genes and the clinical applicability of *in vitro*-derived germ cells for regenerative medicine (Lovell-Badge et al., 2021). To address these challenges, this study aimed to establish an efficient and simplified protocol for differentiating PGC-like cells (PGCLCs) from mouse ESCs. To facilitate PGCLC identification, we generated a Dppa3-mCherry reporter mouse ESC line using CRISPR-Cas9-mediated knock-in. Using these reporter cells, we optimized a differentiation protocol, demonstrating that a culture medium supplemented with ascorbic acid, transferrin, BMP4, and BMP8b induces approximately 50% DPPA3-mCherry-positive PGCLCs by day 5 of differentiation.

## Results

### Generation of Dppa3 reporter mESC through CRISPR Cas9 mediated Knock-in

We generated a Dppa3-mCherry reporter mouse embryonic stem cell (mESC) line, named TDM11, using CRISPR-Cas9-mediated knock-in, as described in the Methods section. Microscopic analysis revealed that TDM11 cells were morphologically identical to the parental TG2A cell line (Figure 1D). In the presence of LIF and serum, approximately 23% of TDM11 cells exhibited mCherry fluorescence (Figure 1E).

**Figure 1:**
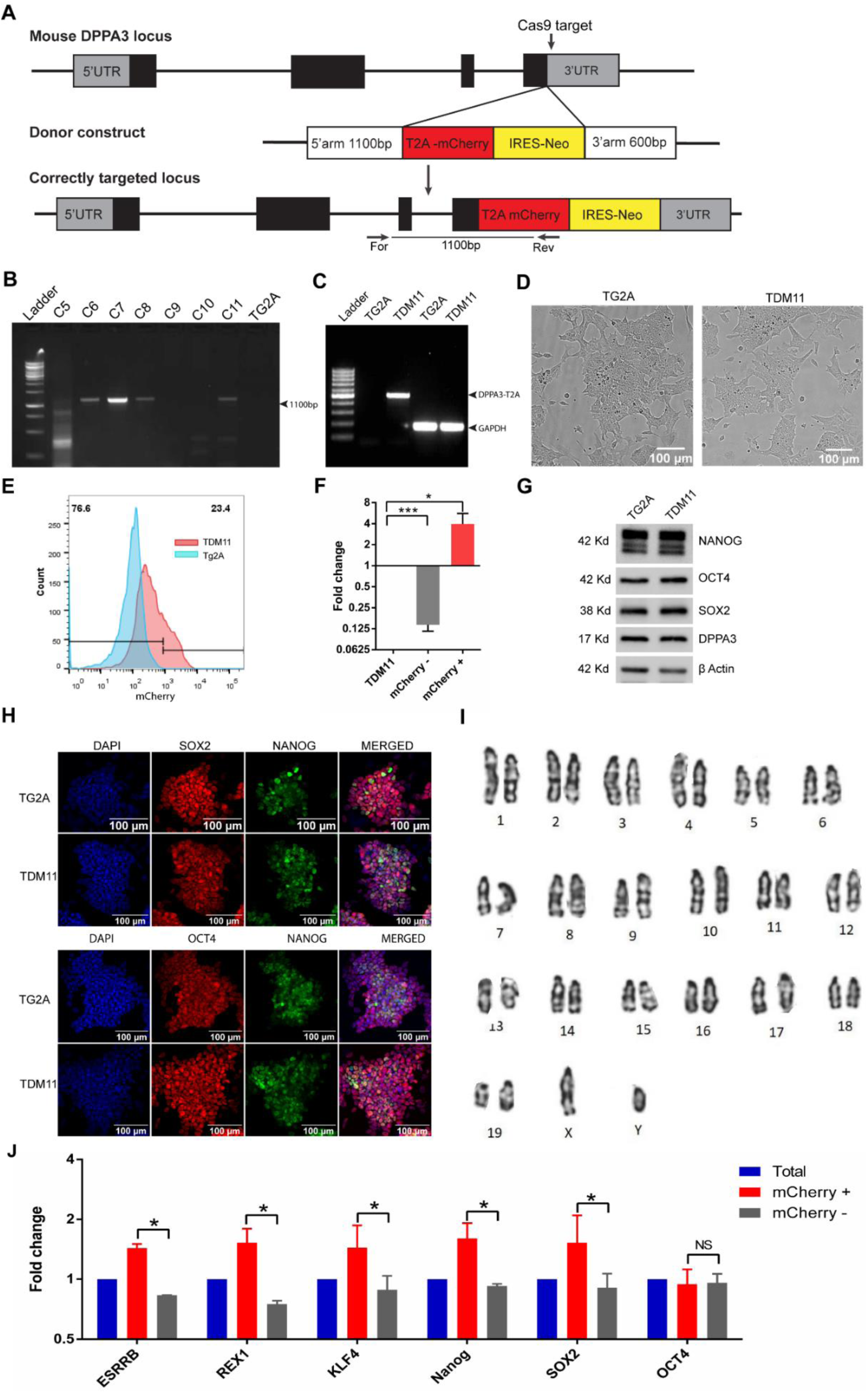
Creation and characterization of TDM11 cell line. A) The schematic representation of targeting vector, specific for Dppa3 last exon and the Knock-in strategy. The For and Res indicates the forward (genome specific) and revers (targeting vector specific) primers used for screening of the correctly integrated clones. B) PCR based screening of neomycin resistant clones for correctly recombination by using forward primer from Dppa3 locus outside of the homologous arm of the targeting vector and second primer specific for T2A-mCherry sequence from the targeting vector. Clone number 11 was named as TDM11. C) The amplification of Dppa3-T2A-mCherry transcript from cDNA of clone 11 (TDM11) using forward primer from first exon of Dppa3 gene and revers primer from T2A sequence. D) Microscopic analysis of colony morphology of TDM11 cells compare to the mother cells TG2A. E) Flowcytometric analysis of mCherry fluorescence of TDM11 cell line grown in LIF plus serum condition compared to TG2A negative cells. F) The qPCR analysis of Dppa3 expression in mCherry positive and negative sorted cells compare to the total population of TDM11 cells. G) The western blotting and H) immunofluorescence analysis for pluripotency factor Oct4, Sox2, Nanog in TDM11 cells compare to the TG2A cells. I) The karyotype of TDM11 cell line.

Fluorescence-activated cell sorting (FACS) of mCherry-positive cells showed a significant upregulation of **Dppa3** expression (4-fold increase), while mCherry-negative cells exhibited a significant downregulation (0.14-fold change) compared to the unsorted TDM11 population (Figure 1F). These results confirm that mCherry fluorescence reliably reports **Dppa3** expression.

Furthermore, the expression levels and profiles of pluripotency markers **Oct4, Sox2,** and **Nanog** in TDM11 cells were identical to those in the parental TG2A cell line (Figures 1G and 1H), confirming the pluripotent nature of TDM11. Additionally, **Dppa3** expression in TDM11 cells were comparable to that in TG2A cells (Figure 1G), indicating that the **T2A-mCherry-IRES-Neo** knock-in at the **Dppa3** locus does not disrupt endogenous **Dppa3** expression.

Karyotype analysis after 10 passages confirmed that TDM11 cells remained euploid (Figure 1I), suggesting genomic stability.

### Dppa3 positive ESCs are poised toward naïve stage of pluripotency

DPPA3 expression was heterogeneous in ESCs, with approximately 23% of cells expressing DPPA3-mCherry. To assess the molecular identity of DPPA3-positive cells in undifferentiated ESCs, we sorted DPPA3-mCherry positive and negative populations and performed qPCR for naïve pluripotency markers. The results showed that DPPA3-mCherry positive cells expressed significantly higher levels of naïve stage markers—Klf4, Rex1, and Esrrb—as well as the pluripotency factors Sox2 and Nanog, compared to DPPA3-mCherry negative cells. In contrast, Oct4 expression was similar in both populations (Figure 1J).

### *In vitro* differentiation of ESCs into PGC like cells (PGCLC)

We differentiated TDM11 cells under embryoid body (EB) culture conditions using basal medium (Table 1) and analyzed the percentage of DPPA3-mCherry-positive cells during differentiation by flow cytometry (Figure S2A). Our results showed that DPPA3 expression declined during the first two days of differentiation, with a subsequent increase beginning on day three (Figure S2B). By day five, approximately 26% of the cells were DPPA3-mCherry positive, suggesting that they had acquired primordial germ cell-like cell (PGCLC) characteristics (Figure S2B).

**Table 1:** Medium composition screened for PGCLC differentiation.

| Basal medium | Basal medium plus Ascorbic acid | Differentiation Medium | Cytokine supplemented differentiation medium |
| --- | --- | --- | --- |
| GMEM, 10% Fetal bovine serum, 0.1mM non-essential amino acid, 0.1mM Mercaptoethanol, 1mM Sodium pyruvate and 1mM GlutaMAX | Basal media + 50 µg/ml Ascorbic acid | Basal media + 50 µg/ml Ascorbic acid + 200 µg/ml Transferrin | Differentiation medium + 100 ng/ml BMP4 + 50 ng/ml BMP8b<br>Or<br>Differentiation medium + 1000 unit/ml LIF |

These findings indicate that PGCLC specification can occur alongside differentiation into the three germ lineages within EB culture conditions. Furthermore, mESCs cultured in LIF plus serum conditions retain the competence to differentiate into PGCLCs *in vitro*.

### Ascorbic acid and transferrin supplementation enhances PGCLC differentiation

We investigated the effect of ascorbic acid (AA) supplementation in basal medium (Table 1) on PGCLC differentiation efficiency. Using our model system, we analyzed the abundance of DPPA3-mCherry-positive cells daily for up to eight days using flow cytometry (Figure S2B). Interestingly, AA supplementation significantly increased the proportion of DPPA3-mCherry-positive cells during differentiation. As expected, during the first two days, the abundance of DPPA3-mCherry-positive cells decreased compared to undifferentiated ESCs. However, beginning on day three, we observed a strong induction of DPPA3-mCherry expression within embryoid bodies (EBs) (Figure S2B). By day five, approximately 34% of the cells in AA-supplemented conditions were DPPA3-mCherry-positive, indicating enhanced PGCLC specification (Figure S2B). Beyond day five, EBs continued to grow, but the proportion of DPPA3-mCherry-positive cells gradually declined each day until day eight.

Furthermore, we examined the effect of transferrin supplementation on PGCLC specification. Notably, adding transferrin to AA-supplemented basal medium—hereafter referred to as differentiation medium (Table 1)—further enhanced differentiation efficiency. By day five, approximately 40% of the cells in EBs were DPPA3-mCherry-positive (Figure S2B), demonstrating an improved induction of PGCLCs. After day five, DPPA3-mCherry expression gradually declined, following a pattern similar to that observed in EBs cultured in basal medium and AA-supplemented medium.

### BMP signalling cascade in PGLCs specifications

To examine that the PGCLC differentiated in our defined system (in differentiation medium) are responsive to these cytokines and if we can enhance the efficiency of PGCLC specification further, we supplemented the recombinant BMP4 and BMP8b in the differentiation medium (Table1). Indeed, we found a significant increase in Dppa3-mCherry positive cells in EBs after supplementation of BMP4 and BMP8b in differentiation media (figure 2B, 2C). At fifth day of differentiation, about 50 % cells were Dppa3-mCherry positive in EBs differentiated into cytokines supplemented differentiation medium, which is significantly high compared to EBs grown in differentiation media lacking the cytokine supplements (Figure 2D). These results clearly showed that PGCLC differentiation in our system are responding to the PGC-specific signalling. These results were further confirmed by Western blotting of Dppa3 at every day of differentiation (**Figure 2E**). Compared to undifferentiated cells, the Dppa3 expression increases significantly on the 2^nd^ day of differentiation and then gradually decrease in EBs differentiated in only differentiation medium, whereas the Dppa3 expression keep increasing up to 4^th^ day of differentiation where the BMP4 and BMP8b is supplemented in differentiation medium (**Figure 2E**). Moreover, we also examined the expression profile of the other germ cell markers; Nanos2, Oct4, Sox2 and Integrin β3. As expected, the expression profile of Nanos2 is like the Dppa3 expression profile, whereas Integrin β3 started expressing at 6 days of differentiation. Integrin β3 expression in EBs differentiated in cytokine supplemented medium is significantly higher than the without cytokine differentiated EBs on 6 days of differentiation (**Figure 2E**).

**Figure 2:**
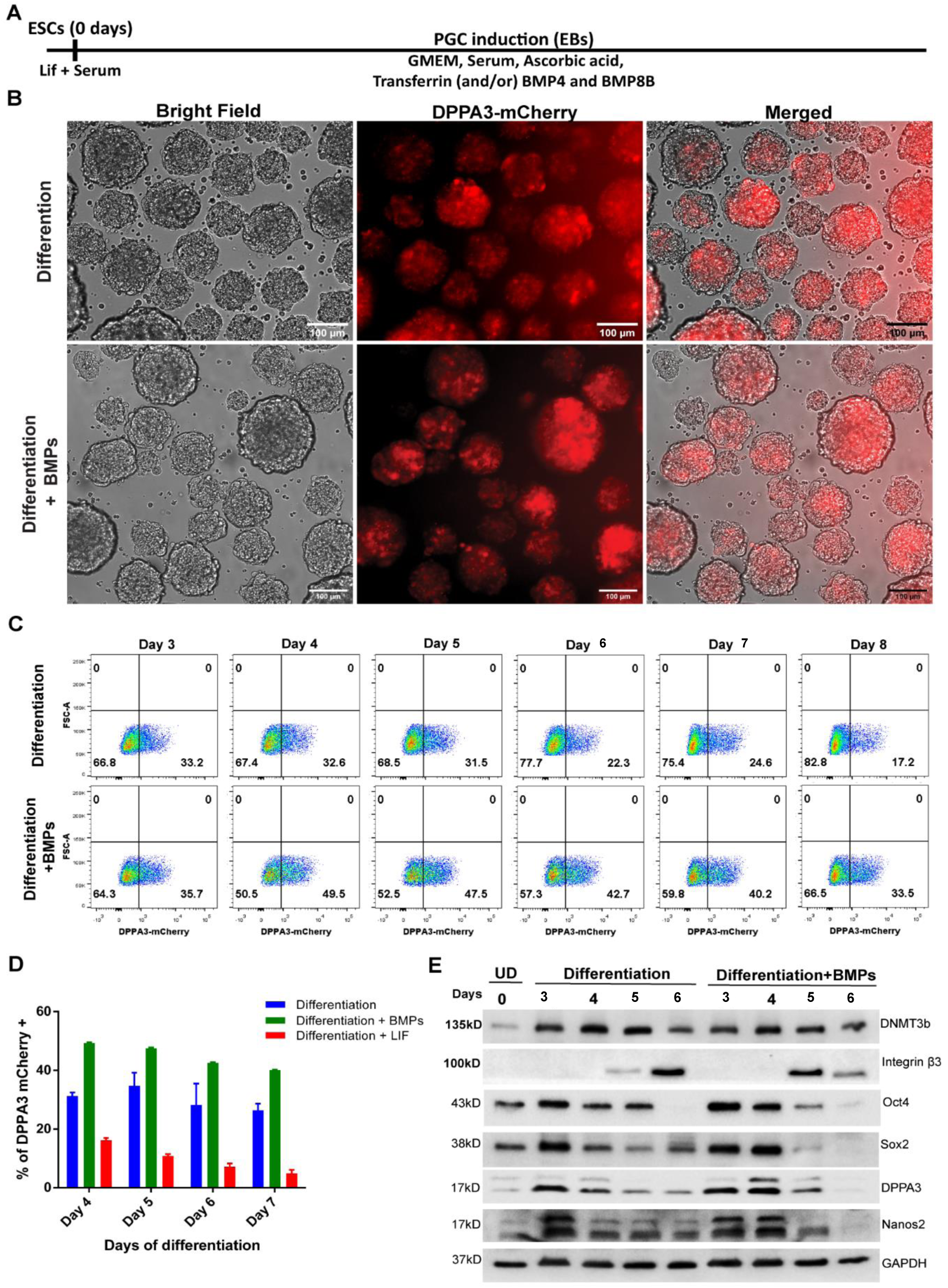
PGCLC induction from mESCs from TDM cells and characterization. A) The schematic strategy for PGCLC induction from mESCs. B) Microscopic analysis Dppa3-mCherry fluorescence and brightfield of the fourth day differentiated EBs at fourth day of differentiation, differentiated in differentiation media and differentiation media supplemented with BMP4 and BMP8b **C**). Flowcytometric analysis of Dppa3-mCherry positive cells from 3^rd^ day of differentiation to 8^th^ day of differentiation. D) The percentage of the Dppa3-mCherry positive cells in EBs differentiated in cytokine supplemented differentiation medium compared to only in differentiation medium. E) Western blot analysis of the PGC expressed genes during differentiation compared to undifferentiated (UD) TDM11 cells at day 0.

### Effects of LIF in PGLC specifications

To investigate the role of LIF (Leukemia Inhibitory Factor) in the specification of primordial germ cell-like cells (PGCLCs) within our differentiation system, we added LIF to the differentiation medium, as detailed in Table 1. We observed that the percentage of Dppa3-mCherry positive cells decreased steadily from the second to the seventh day of differentiation. By the fifth day, only around 15% of the cells were Dppa3-mCherry positive, which is roughly three times lower than embryoid bodies (EBs) cultured without LIF in the differentiation medium (Figure 2D). LIF is known to maintain embryonic stem cells (ESCs) in an undifferentiated state. These findings suggest that LIF inhibits PGC specification by preventing ESCs from differentiating in our protocol.

### Molecular characterization of derived PGCLC

To examine the molecular characteristics of Dppa3-mCherry positive cells within the EBs, which were differentiated in both standard and BMP supplemented media, we sorted the Dppa3-mCherry positive and negative cells. We analyzed the expression of PGC-related genes in comparison to undifferentiated cells. The pluripotency markers Oct4, Sox2, and Nanog were expressed equally in Dppa3-mCherry positive PGCLCs and undifferentiated ESCs (Figure 3B). As expected, Sox2 and Nanog were significantly downregulated in the negative differentiated cells (Figure 3B), although Oct4 remained equally expressed in both positive and negative cells, possibly due to its presence in other cell lineages.

**Figure 3:**
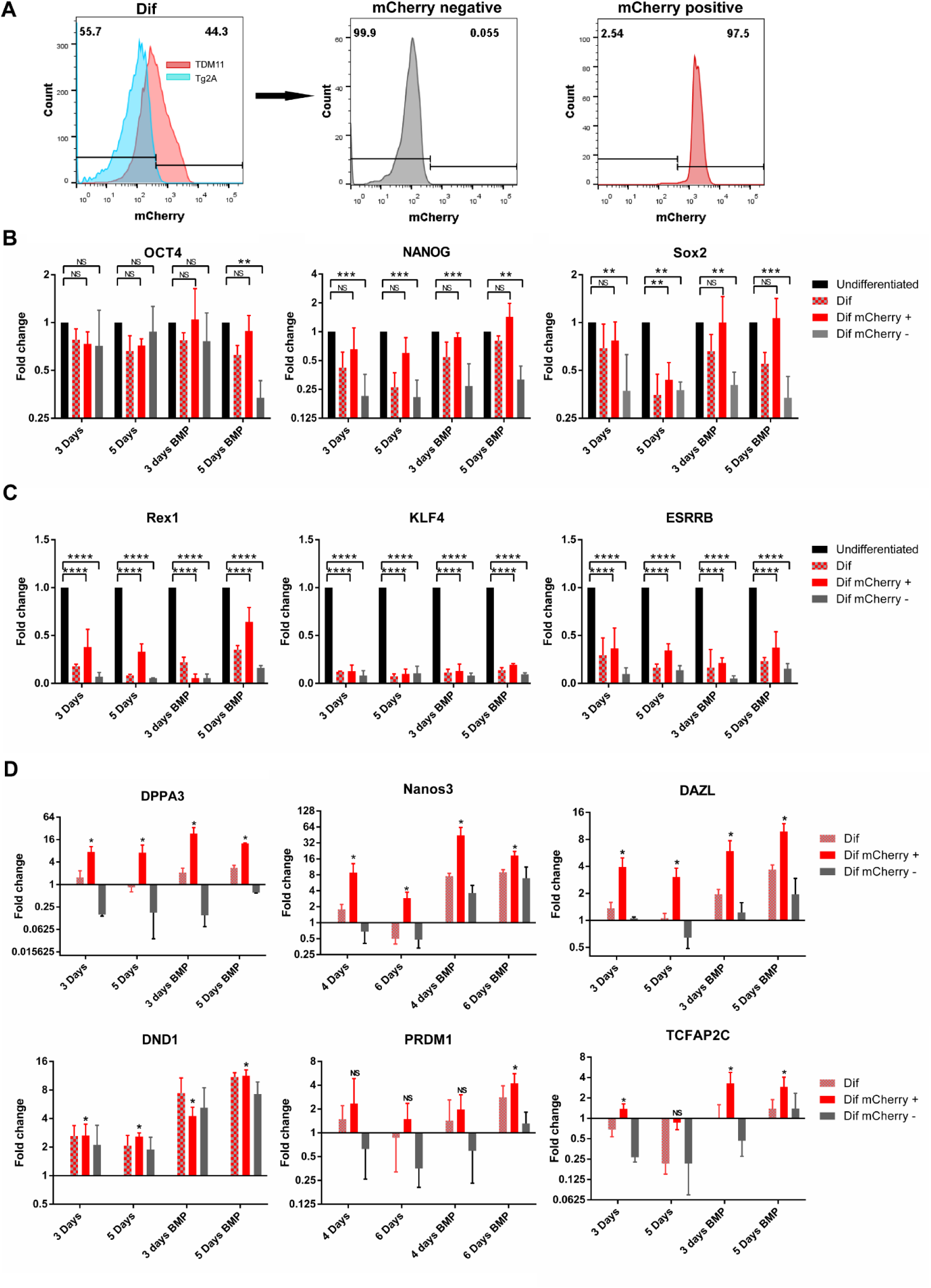
Molecular identity of invitro differentiated PGCLCs. A) Representative image of flowcytometric analysis of Dppa3-mCherry positive and negative cell sorting from differentiating EBs. B) The expression of PGC expressed pluripotency factors in Dppa3-mCherry positive and negative sorted cell, from the EBs differentiated in differentiation media compared to undifferentiated ESCs. C) The expression of naïve stage of pluripotency factors in Dppa3-mCherry positive and negative cell sorted cells from differentiating EBs compared to undifferentiated ESCs. D) The expression of PGC expressed gene in Dppa3-mCherry positive and negative cell sorted cells from differentiating EBs compared to undifferentiated ESCs.

Since Dppa3 is also expressed in undifferentiated ESCs at the naïve stage (as shown in Figure 1H), we evaluated naïve state markers Klf4, Rex1, and Esrrb to ensure that the Dppa3-mCherry positive cells in differentiating EBs are not naïve stage cells. All three markers were significantly downregulated in both Dppa3-mCherry positive and negative cells compared to undifferentiated ESCs, as presented in Figure 3C.

We further assessed the expression of PGC-specific genes like Dppa3, Dazl, and Nanos3, which were significantly enriched in Dppa3-mCherry positive cells in comparison to both undifferentiated and negative cells (Figure 3D). Dnd1, another PGC-expressed gene, showed marked upregulation in positive Dppa3-mCherry cells compared to undifferentiated cells, although it was also expressed in negative cells (Figure 3D). The expression levels of Prdm1 and Tcfap2C in Dppa3-mCherry positive cells were similar to those in undifferentiated cells (Figure 3D), reinforcing our conclusion that Dppa3-mCherry positive cells in differentiating EBs are indeed PGCLCs.

### Functional characterization of PGCLCs

Cell cycle analysis revealed that Dppa3-mCherry positive cells, sorted by FACS on the fifth day of differentiation, were enriched in the G2 phase (42%) compared to the negative cells at the same stage (22%) (Figure 4A). This suggests that Dppa3-mCherry positive PGCLCs proliferate more slowly after induction in EBs, while negative cells continue to proliferate. This also explains the reduced abundance of Dppa3-mCherry positive cells after the fifth day of differentiation (Figure 2C).

**Figure 4:**
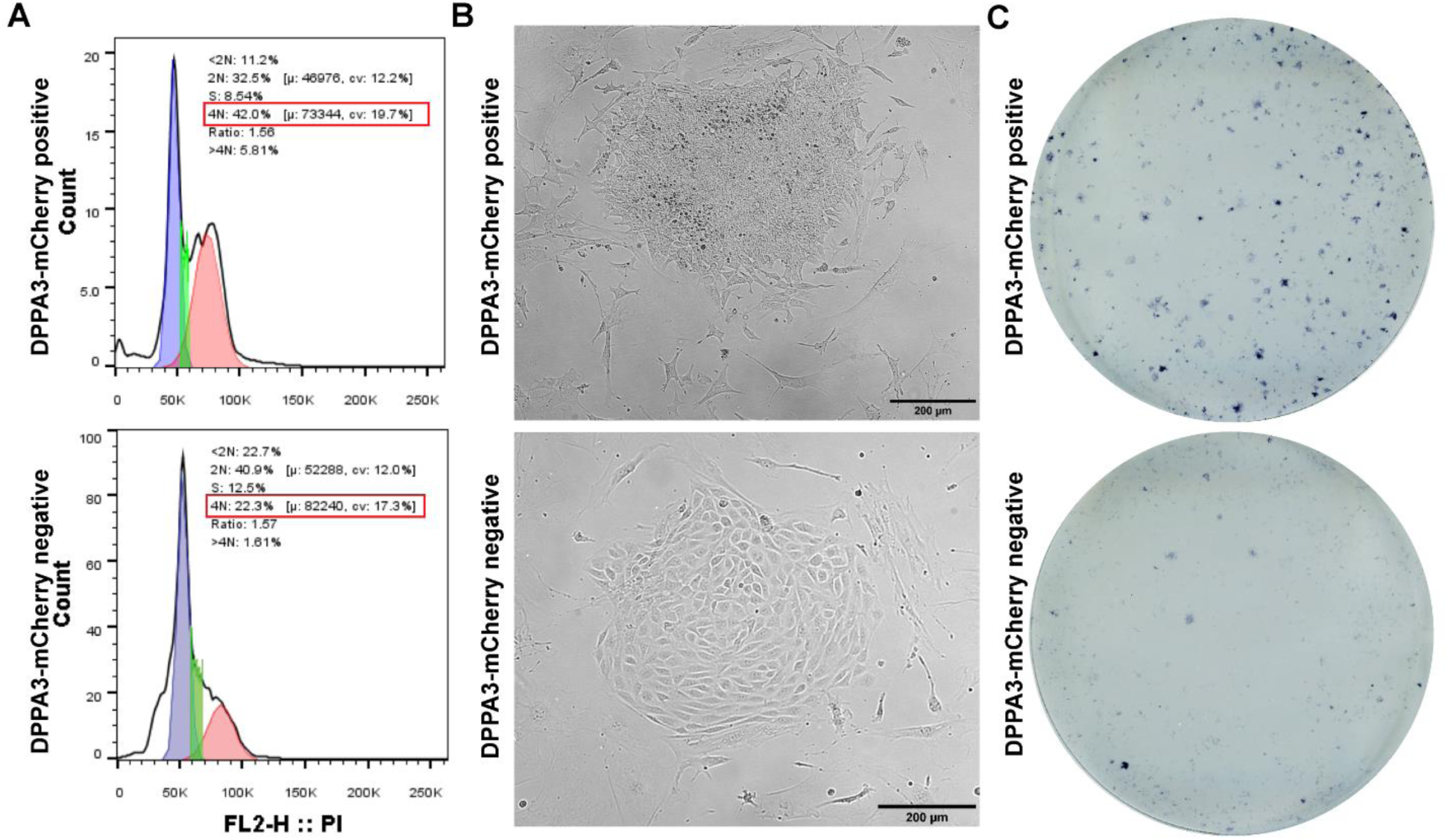
Functional characterization of PGCLC. A) Cell cycle assay of Dppa3-mCherry positive and negative sorted PGCLC at 4^th^ day EBs differentiated in differentiation media. B) Morphology of colonies developed from Dppa3-mCherry positive and negative sorted PGCLC, grown in EGC medium. C) Alkaline phosphatase staining of the colonies developed from Dppa3-mCherry positive and negative sorted PGCLC, grown in EGC media.

For further functional assessment, we cultured sorted Dppa3-mCherry positive and negative cells from EBs on mouse embryonic fibroblasts (MEFs) using embryonic germ cell (EGC) media, as described by (Vincent et al., 2011). In the presence of retinoic acid (a component of EGC media), PGCs form alkaline phosphatase-positive colonies, whereas undifferentiated ESCs differentiate into various somatic lineages (Dyce et al., 2018; Zhang et al., 2015). Dppa3-mCherry positive sorted PGCLCs from the fifth day of differentiation formed stem cell-like colonies, while the negative cells differentiated into flat cuboidal cells (Figure 4B). Moreover, colonies derived from positive PGCLCs exhibited strong alkaline phosphatase staining, contrary to the unstained colonies from negative cells (Figure 4C). We also established continuously growing EGC cell lines from colonies developed from positive PGCLCs in EGC medium. These findings confirm that the derived PGCLCs functionally resemble in vivo PGCs.

## Discussion

Primordial germ cell (PGC) specification in mammals, including mice and humans, involves three key processes: suppression of somatic genes, regaining potential pluripotency and latent totipotency, and extensive epigenetic reprogramming. These processes are driven by cytokine signaling. Recent studies have successfully recreated PGC pathways *in vitro* from pluripotent stem cells, using temporal manipulation of signaling pathways through cytokines and growth factors (Hayashi et al., 2011; Hikabe et al., 2016; Nakaki et al., 2013). When *in vitro*-derived PGC-like cells (PGCLCs) are transplanted into neonatal testes, they can differentiate into functional gametes (Sosa et al., 2018).

We have developed an efficient method to differentiate PGCLCs from mouse embryonic stem cells (ESCs) using ascorbic acid (vitamin C) supplementation. A novel Dppa3-mCherry reporter mouse ESC line was created by inserting a T2A-mCherry-IRES-Neo cassette into the Dppa3 gene locus. This TDM11 cell line accurately reflects DPPA3 gene expression. Approximately 23% of ESCs in LIF plus serum medium express the DPPA3 gene, consistent with previous studies (Hayashi et al., 2008).

Embryoid bodies (EBs), which are 3D structures formed by aggregating ESCs, can mimic early embryonic development and contain PGCs along with other cell lineages. In our experiments, about 25% of cells in 5-day EBs were DPPA3-mCherry positive. During PGC development, extensive epigenetic reprogramming occurs, facilitated by enzymes like the Ten-eleven translocation (Tet) enzymes, which are involved in DNA demethylation (Guo et al., 2017; Miyoshi et al., 2016; Seisenberger et al., 2012; Sekl et al., 2007). (Wu & Zhang, 2011; Yamaguchi et al., 2013). The Prdm14, an early PGC expression factor, promotes DNA demethylation through Tet enzyme activation (Okashita et al., 2016). Ascorbic acid acts as a cofactor for Tet enzymes, enhancing germ cell development (DiTroia et al., 2019c). Recently, it was demonstrated that the maternal ascorbic acid plays important role in germ cell development, and maternal deficiency leads to reduce the number of germ cells (DiTroia et al., 2019b). Moreover, AA directly activates Tet enzymes in ESCs and leads to activate the expression of the germ cell-specific genes such as Dazl (Blaschke et al., 2013). Similarly, Transferrin, secreted by Sertoli cells, supports PGC self-renewal, as indicated in chickens and possibly applicable in mice (Gelly et al., 1994; Whyte et al., 2015).

We examined the effects of ascorbic acid and transferrin on EBs-mediated PGC differentiation by monitoring Dppa3-mCherry expression. Ascorbic acid significantly increased the number of DPPA3-mCherry positive cells (Figure S2). BMP4 and BMP8b, critical for *in vivo* PGC specification, further enhanced the abundance of these cells when added (Figure 2D). DPPA3-mCherry positive cells in ascorbic acid-supplemented EBs expressed important PGC markers such as Nanos3, DazL, and Dnd1, and retained pluripotency factors like Oct4, Nanog, and Sox2 (Figure 3B and 3D). These PGCLCs displayed characteristics typical of PGCs, such as slow growth and quiescence in the G2 phase of the cell cycle (**Figure 3B**). The inner cells mass (ICM) markers, Rex1, Klf4, and Esrrb are significantly downregulated Dppa3-mCherry positive cells PGCLC compared to undifferentiated ESCs, while retaining the expression of Prdm14, and Tfap2C, which are common factors between ESCs and PGCs (**Figure 3C and 3D**) confirm the identity of PGCLCs.

A key functional characteristic of PGCs is their slow cell growth and their quiescence nature at G2 phase of cell cycle (Sekl et al., 2007; Takehara & Matsui, 2019). We found that DPPA3-mCherry positive PGCLC enriched in G2 phase of cell cycle (figure 4A). One of the functional properties of PGC, which can distinguish between PGC, ESC and differentiated cells is responsiveness to the retinoic acid (RA), whereas it actively stimulates the proliferation of PGCs while induce rapid differentiation in ESCs (Dyce et al., 2018; Zhang et al., 2015). We found that the sorted Dppa3-mCherry positive cells from developed as alkaline phosphatase positive colonies in the presence of RA while Dppa3-mCherry negative differentiated cells dose not stains for alkaline phosphatase (**Figure 4B and 4C**). Furthermore, we derived the continuously growing embryonic germ cell lines (EGC) by isolating individual RA resistant colonies, developed from Dppa3-mCherry positive cells, and grown in LIF containing ESCs growth media, corroborates with previous studies (Leitch et al., 2010; Shamblott et al., 1998). These results further demonstrate the functionality of differentiated PGCLC in our system.

In summary, we have developed an efficient and simple PGCLC differentiation protocol from mESCs. This protocol will be useful not only to differentiate the functional gamete but open the possibility to characterize the role and molecular function of novel genes in PGC specification, *in vitro*.

## Material and methods

### Creation of TDM11 cell line and culture

The mESCs E14-TG2A (TG2A) were grown in LIF + serum medium condition (GMEM, 10% fetal bovine serum, 0.1mM non-essential amino acid, 1000 units/ml LIF, 0.1mM 2-Mercaptoethanol, 1mM Sodium pyruvate) on gelatine coted cell culture plates. To quantify the live PGCLC population by faithful reporting of Dppa3 gene, we knocked-in T2A-mCherry-IRES-Neo cassette at Dppa3 locus in frame, right before the stop codon, using CRISPR-Cas9 mediated gene editing as depicted in **figure 1A**. The targeting plasmid is shown in figure 1A. The twin guides RNA oligos, specific for Dppa3 targeted region was cloned under the regulation of U6 promoter in another plasmid which also have SpCas9 cloned under regulation chicken β-actin promoter. Both the targeting plasmid and twin Guide-RNA plus SpCas9 vector was co-nucleofected into TG2A cells. The positively integrated cells were selected by adding 150 µg/ml neomycin in the growth medium for seven days. The individual resistant colonies were picked up under microscope and dissociated with TrypLE (Invitrogen, USA). Neomycin resistant clones were screened by PCR-based genotyping for recombination of targeting cassette at Dppa3 locus with an expected amplificon size of about 1.1 kb **(figure 1B)**. One of the positive clones, clone number 11 (named TDM11) were further confirmed by amplification of Dppa3-T2A-mCherry recombination-specific transcript from cDNA (**figure 1C)**. To confirm that the integration is in the frame of Dppa3 gene and T2A-mCherry start after last codon of Dppa3 transcript, the amplicon was sequenced (**figure S1A**). The dissociated cells from individual colonies were expanded and genotyping for the positive integration was done using integration specific PCR. The one of the positive clones (C11) was further characterized and named as TDM11 (TG2A Dppa3 mCherry clone 11).

### PGCLC induction from TDM11 cell line

TDM11 cells, grown in LIF plus serum condition were dissociated from plate and about 500000 cells were seeded in nonadherent culture P60 plates in three ml of different media compositions as shown in table 1. Up to second day of differentiation 1 ml of media added per day and afterward half of the medium from differentiation plates was replenished with new medium. The Ebs from plates were harvested and washed with PBS and dissociated into single cell by treating them in TrypLE for five minutes for flowcytometric analysis.

### Derivation of EGC from PGCLCs

After growing EBs in differentiation medium for five days, the cells were dissociated into single cell suspension and Dppa3-mCherry positive and negative cells were sorted by FACS. About 3000 cells were seeded on inactivated mouse embryonic fibroblasts (MEFs) in EGC medium (GMEM, 10% FBS, 1mM Sodium Pyruvate, 0.1mM Nonessential Amino Acid, 1mM GlutaMax, 0.1mM 2-mMercaptoethanol, 2µM Retinoic Acid, LIF, 30 ng/ml SCF, 15 ng/ml bFGF2). The medium was replenished every day and after five days of growth, the grown colonies either harvested or assayed for alkaline phosphatase activity. The harvested cells were further propagated in ESCs culture media.

### Cell cycle analysis

The cells were sorted from 5^th^ day differentiated EBs. The sorted cells from differentiating EBs were fixed using 70% ice cold ethanol and stained with Propidium Iodide. The stained cells were treated with RNase to remove RNA for 15 minutes and analysed using flow-cytometer. The cell cycle was plotted using FlowJo software.

## Acknowledgements

K.T. was supported by funding from Department of Biotechnology, Ministry of Science and technology, Government of India, and CSIR Bhatnagar Fellowship (CSIRHRD/BFS2024/03/01) from Council of Scientific and Industrial Research, Ministry of Science and technology, Government of India.

---

**Figure S1.**
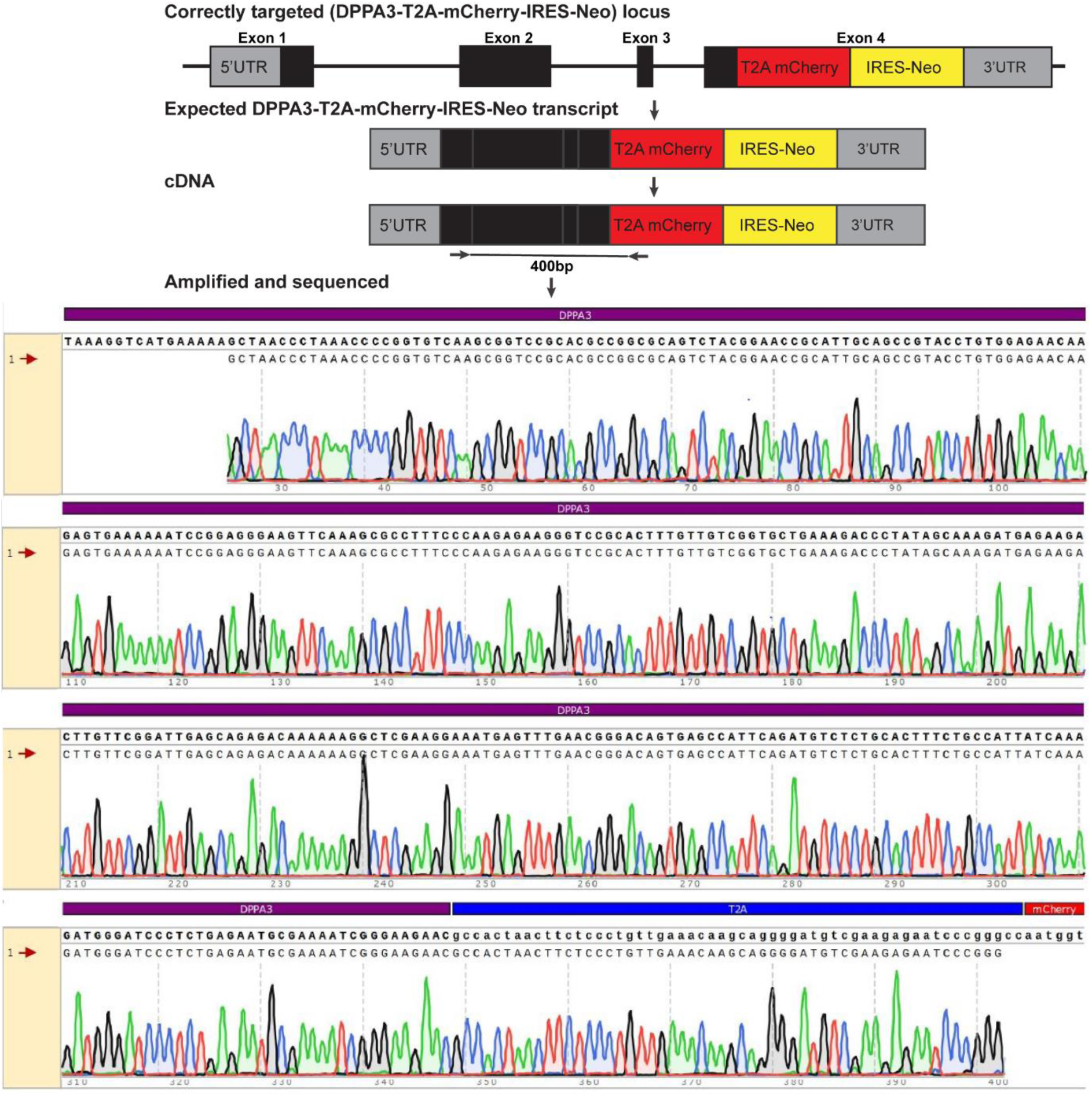
Sanger sequencing of Dppa3-mCherry transcript, isolated and revers transcribed from TDM11 cells.

**Figure S2:**
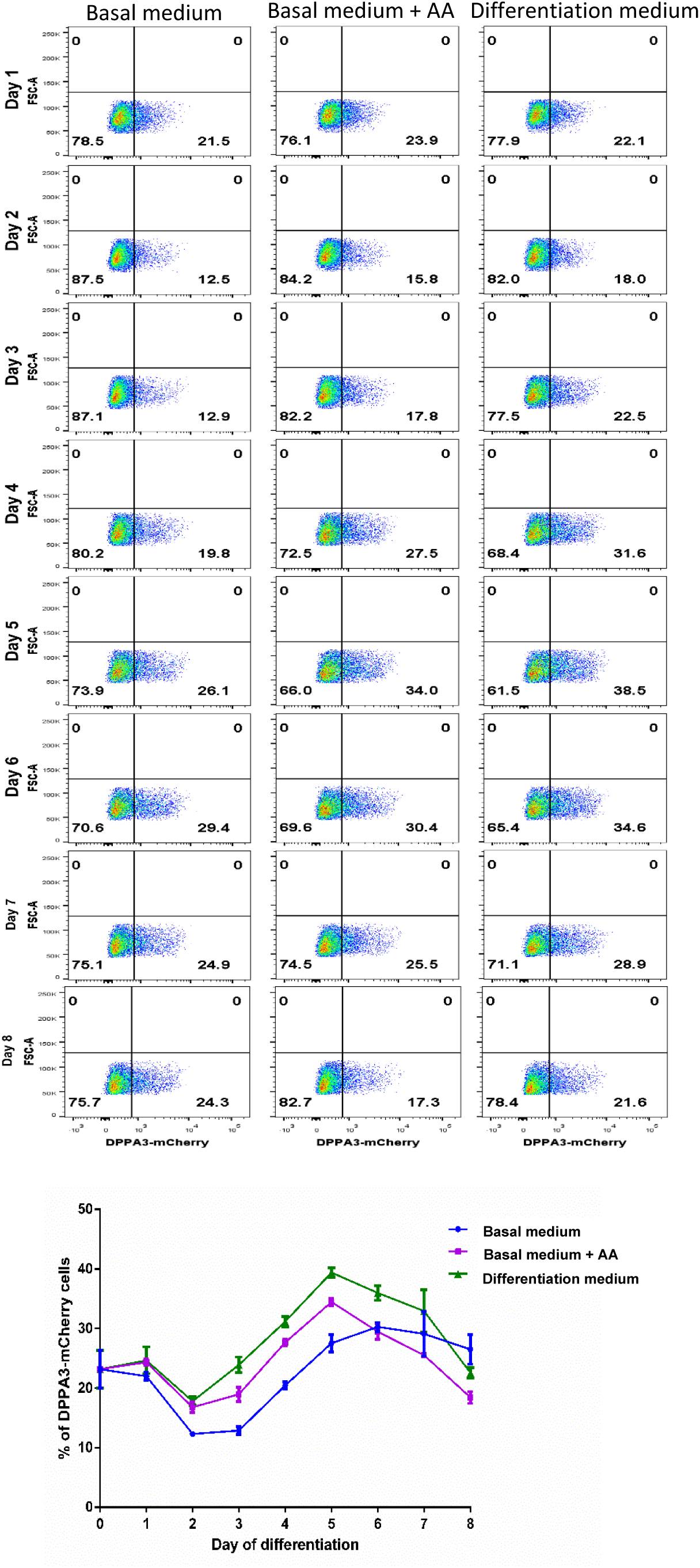
Flowcytometric analysis of DPPA3-mCherry positive cells during differentiation. Flowcytometric analysis of differentiated cells, differentiated in Basal medium, Basal medium plus Ascorbic Acid (AA) and in differentiation medium, up to 8^th^ day of differentiation. The data of the percentage of Dppa3-mCherry positive cells during differentiation in Basal medium, Basal Medium plus AA and Different medium was plotted.

## References

Blaschke, K., Ebata, K. T., Karimi, M. M., Zepeda-Martínez, J. A., Goyal, P., Mahapatra, S., Tam, A., Laird, D. J., Hirst, M., Rao, A., Lorincz, M. C., & Ramalho-Santos, M. (2013). Vitamin C induces Tet-dependent DNA demethylation and a blastocyst-like state in ES cells. Nature, 500(7461), 222–226. 10.1038/nature12362

Conley, B. J., Trounson, A. O., & Mollard, R. (2004). Human embryonic stem cells form embryoid bodies containing visceral endoderm-like derivatives. Fetal Diagnosis and Therapy, 19(3), 218–223. 10.1159/000076701

de Sousa Lopes, S. M. C., Hayashi, K., & Surani, M. A. (2007). Proximal visceral endoderm and extraembryonic ectoderm regulate the formation of primordial germ cell precursors. BMC Developmental Biology, 7. 10.1186/1471-213X-7-140

DiTroia, S. P., Percharde, M., Guerquin, M. J., Wall, E., Collignon, E., Ebata, K. T., Mesh, K., Mahesula, S., Agathocleous, M., Laird, D. J., Livera, G., & Ramalho-Santos, M. (2019a). Maternal vitamin C regulates reprogramming of DNA methylation and germline development. Nature, 573(7773), 271–275. 10.1038/s41586-019-1536-1

Dyce, P. W., Tenn, N., & Kidder, G. M. (2018). Retinoic acid enhances germ cell differentiation of mouse skin-derived stem cells. Journal of Ovarian Research, 11(1). 10.1186/s13048-018-0390-3

Gelly, J.-L., Richoux, J.-P., & Grignon, G. (1994). Cell Tissue Research Immunolocalization of albumin and transferrin in germ and Sertoli cells during rat gonadal morphogenesis and postnatal development of the testis cells. In Cell Tissue Res (Vol. 276).

Guo, H., Hu, B., Yan, L., Yong, J., Wu, Y., Gao, Y., Guo, F., Hou, Y., Fan, X., Dong, J., Wang, X., Zhu, X., Yan, J., Wei, Y., Jin, H., Zhang, W., Wen, L., Tang, F., & Qiao, J. (2017). DNA methylation and chromatin accessibility profiling of mouse and human fetal germ cells. Cell Research, 27(2), 165–183. 10.1038/cr.2016.128

Hayashi, K., Lopes, S. M. C. de S., Tang, F., & Surani, M. A. (2008). Dynamic Equilibrium and Heterogeneity of Mouse Pluripotent Stem Cells with Distinct Functional and Epigenetic States. Cell Stem Cell, 3(4), 391–401. 10.1016/j.stem.2008.07.027

Hayashi, K., Ohta, H., Kurimoto, K., Aramaki, S., & Saitou, M. (2011). Reconstitution of the mouse germ cell specification pathway in culture by pluripotent stem cells. Cell, 146(4), 519–532. 10.1016/j.cell.2011.06.052

Hikabe, O., Hamazaki, N., Nagamatsu, G., Obata, Y., Hirao, Y., Hamada, N., Shimamoto, S., Imamura, T., Nakashima, K., Saitou, M., & Hayashi, K. (2016). Reconstitution in vitro of the entire cycle of the mouse female germ line. Nature, 539(7628), 299–303. 10.1038/nature20104

Irie, N., Weinberger, L., Tang, W. W. C., Kobayashi, T., Viukov, S., Manor, Y. S., Dietmann, S., Hanna, J. H., & Surani, M. A. (2015). SOX17 is a critical specifier of human primordial germ cell fate. Cell, 160(1–2), 253–268. 10.1016/j.cell.2014.12.013

Kurimoto, K., Yamaji, M., Seki, Y., & Saitou, M. (2008). Specification of the germ cell lineage in mice: A process orchestrated by the PR-domain proteins, Blimp1 and Prdm14. In Cell Cycle (Vol. 7, Issue 22, pp. 3514–3518). Taylor and Francis Inc. 10.4161/cc.7.22.6979

Leitch, H. G., Blair, K., Mansfield, W., Ayetey, H., Humphreys, P., Nichols, J., Surani, M. A., & Smith, A. (2010). Embryonic germ cells from mice and rats exhibit properties consistent with a generic pluripotent ground state. Development, 137(14), 2279–2287. 10.1242/dev.050427

Lovell-Badge, R., Anthony, E., Barker, R. A., Bubela, T., Brivanlou, A. H., Carpenter, M., Charo, R. A., Clark, A., Clayton, E., Cong, Y., Daley, G. Q., Fu, J., Fujita, M., Greenfield, A., Goldman, S. A., Hill, L., Hyun, I., Isasi, R., Kahn, J., … Zhai, X. (2021). ISSCR Guidelines for Stem Cell Research and Clinical Translation: The 2021 update. In Stem Cell Reports (Vol. 16, Issue 6, pp. 1398–1408). Cell Press. 10.1016/j.stemcr.2021.05.012

Miyoshi, N., Stel, J. M., Shioda, K., Qu, N., Odahima, J., Mitsunaga, S., Zhang, X., Nagano, M., Hochedlinger, K., Isselbacher, K. J., & Shioda, T. (2016). Erasure of DNA methylation, genomic imprints, and epimutations in a primordial germ-cell model derived from mouse pluripotent stem cells. Proceedings of the National Academy of Sciences of the United States of America, 113(34), 9545–9550. 10.1073/pnas.1610259113

Nakaki, F., Hayashi, K., Ohta, H., Kurimoto, K., Yabuta, Y., & Saitou, M. (2013). Induction of mouse germ-cell fate by transcription factors in vitro. Nature, 501(7466), 222–226. 10.1038/nature12417

Ohinata, Y., Sano, M., Shigeta, M., Yamanaka, K., & Saitou, M. (2008). A comprehensive, non-invasive visualization of primordial germ cell development in mice by the Prdm1-mVenus and Dppa3-ECFP double transgenic reporter. Reproduction, 136(4), 503–514. 10.1530/REP-08-0053

Okashita, N., Suwa, Y., Nishimura, O., Sakashita, N., Kadota, M., Nagamatsu, G., Kawaguchi, M., Kashida, H., Nakajima, A., Tachibana, M., & Seki, Y. (2016). PRDM14 Drives OCT3/4 Recruitment via Active Demethylation in the Transition from Primed to Naive Pluripotency. Stem Cell Reports, 7(6), 1072–1086. 10.1016/j.stemcr.2016.10.007

Seisenberger, S., Andrews, S., Krueger, F., Arand, J., Walter, J., Santos, F., Popp, C., Thienpont, B., Dean, W., & Reik, W. (2012). The Dynamics of Genome-wide DNA Methylation Reprogramming in Mouse Primordial Germ Cells. Molecular Cell, 48(6), 849–862. 10.1016/j.molcel.2012.11.001

Sekl, Y., Yamaji, M., Yabuta, Y., Sano, M., Shigeta, M., Matsui, Y., Saga, Y., Tachibana, M., Shinkai, Y., & Saitou, M. (2007). Cellular dynamics associated with the genome-wide epigenetic reprogramming in migrating primordial germ cells in mice. Development, 134(14), 2627–2638. 10.1242/dev.005611

Shamblott, M. J., Axelman, J., Wang, S., Bugg, E. M., Littlefield, J. W., Donovan, P. J., Blumenthal, P. D., Huggins, G. R., & Gearhart, J. D. (1998). Derivation of pluripotent stem cells from cultured human primordial germ cells (alkaline phosphataseembryoid bodyembryonic stem cellembryonic germ cell). In Developmental Biology (Vol. 95). www.pnas.org.

Sharlip, I. D., Jarow, J. P., Belker, A. M., Lipshultz, L. I., Sigman, M., Thomas, A. J., Schlegel, P. N., Howards, S. S., Nehra, A., Damewood, M. D., Overstreet, J. W., & Sadovsky, R. (2002). Best practice policies for male infertility.

Smela, M. P., Sybirna, A., Wong, F. C. K., & Azim Surani, M. (2019). Testing the role of sox15 in human primordial germ cell fate [version 2; peer review: 2 approved]. Wellcome Open Research, 4. 10.12688/wellcomeopenres.15381.1

Sosa, E., Chen, D., Rojas, E. J., Hennebold, J. D., Peters, K. A., Wu, Z., Lam, T. N., Mitchell, J. M., Sukhwani, M., Tailor, R. C., Meistrich, M. L., Orwig, K. E., Shetty, G., & Clark, A. T. (2018). Differentiation of primate primordial germ cell-like cells following transplantation into the adult gonadal niche. Nature Communications, 9(1). 10.1038/s41467-018-07740-7

Sugawa, F., Araúzo-Bravo, M. J., Yoon, J., Kim, K., Aramaki, S., Wu, G., Stehling, M., Psathaki, O. E., Hübner, K., & Schöler, H. R. (2015). Human primordial germ cell commitment in vitro associates with a unique PRDM14 expression profile . The EMBO Journal, 34(8), 1009–1024. 10.15252/embj.201488049

Takehara, A., & Matsui, Y. (2019). Shortened G1 phase of cell cycle and decreased histone H3K27 methylation are associated with AKT-induced enhancement of primordial germ cell reprogramming. Development Growth and Differentiation, 61(6), 357–364. 10.1111/dgd.12621

Vincent, J. J., Li, Z., Lee, S. A., Liu, X., Etter, M. O., Diaz-Perez, S. v., Taylor, S. K., Gkountela, S., Lindgren, A. G., & Clark, A. T. (2011). Single cell analysis facilitates staging of blimp1-dependent primordial germ cells derived from mouse embryonic stem cells. PLoS ONE, 6(12). 10.1371/journal.pone.0028960

Whyte, J., Glover, J. D., Woodcock, M., Brzeszczynska, J., Taylor, L., Sherman, A., Kaiser, P., & McGrew, M. J. (2015). FGF, Insulin, and SMAD Signaling Cooperate for Avian Primordial Germ Cell Self-Renewal. Stem Cell Reports, 5(6), 1171–1182. 10.1016/j.stemcr.2015.10.008

Wu, H., & Zhang, Y. (2011). Mechanisms and functions of Tet proteinmediated 5-methylcytosine oxidation. In Genes and Development (Vol. 25, Issue 23, pp. 2436–2452). 10.1101/gad.179184.111

Yamaguchi, S., Shen, L., Liu, Y., Sendler, D., & Zhang, Y. (2013). Role of Tet1 in erasure of genomic imprinting. Nature, 504(7480), 460–464. 10.1038/nature12805

Yamashiro, C., Sasaki, K., Yokobayashi, S., Kojima, Y., & Saitou, M. (2020). Generation of human oogonia from induced pluripotent stem cells in culture. Nature Protocols, 15(4), 1560–1583. 10.1038/s41596-020-0297-5

Ying, Y., Qi, X., & Zhao, G.-Q. (2001). Induction of primordial germ cells from murine epiblasts by synergistic action of BMP4 and BMP8B signaling pathways. In PNAS July (Vol. 3, Issue 14). www.pnas.orgcgidoi10.1073pnas.151242798

Zhang, J., Gao, Y., Yu, M., Wu, H., Ai, Z., Wu, Y., Liu, H., Du, J., Guo, Z., & Zhang, Y. (2015). Retinoic acid induces embryonic stem cell differentiation by altering both encoding RNA and microRNA expression. PLoS ONE, 10(7). 10.1371/journal.pone.0132566

